# Parallel patterns of development between independent cases of hybrid seed inviability in *Mimulus*

**DOI:** 10.1101/458752

**Authors:** Jenn M. Coughlan, John H. Willis

**Affiliations:** Biological Sciences, Duke University 25 Science Drive, Durham NC; Biology Department, University of North Carolina, Chapel Hill

**Keywords:** Cryptic Species, Endosperm, Parental Conflict, Reproductive Isolation, Parallel Evolution

## Abstract

**Rationale:** Hybrid seed inviability (HSI) is a common reproductive barrier in angiosperms, yet the evolutionary and developmental drivers of HSI remain largely unknown. We test whether conflict between maternal and paternal interests in resource allocation to developing offspring (i.e. parental conflict) are associated with HSI and determine the degree of developmental parallelism between independent incidences of HSI in *Mimulus*.

**Methods:** We quantified HSI between *M. guttatus* and two clades of *M. decorus* with oppositely asymmetric incompatibilities and surveyed development of hybrid and parental seeds.

**Key Results:** Crosses between *M. guttatus* and both clades of *M. decorus* show parent-of-origin effects on reciprocal F1 seed development, but in opposing directions. Inviable hybrid seeds exhibit paternal excess phenotypes, wherein endosperm is large and chaotic while viable hybrid seeds produce endosperm cells that are smaller and less prolific (i.e. maternal-excess phenotypes).

**Main Conclusions:** We find strong parent-of-origin effects on development in reciprocal F1s in multiple incidences of HSI in *Mimulus*. These patterns suggest that parental conflict may be an important force generating HSI in this group, and mismatches between maternal and paternal contributions to developing seeds result in repeatable developmental defects in hybrids.

## Introduction

The repeatability of speciation is a contentious topic in evolutionary biology. Many pre-zygotic barriers can be attributed to adaptations to the abiotic or biotic environment (Coyne & Orr 2004), and the repeatability of extrinsic reproductive isolation may be directly related to the repeatability of adaptation (i.e. Rundle *et al*. 2000; Rolán-Alvarez *et al*. 2004; Stankowski 2013; Soria-Carrasco *et al*. 2014; Stuart *et al*. 2017, though see Lowry *et al*. 2008). Yet intrinsic post-zygotic reproductive isolation can also manifest canonical hybrid phenotypes (e.g. CMS hybrid sterility in plants-reviewed in Hanson & Bentolila 2004; Chase 2007; Rieseberg & Blackman 2010; hybrid necrosis in plant and fungi-reviewed in Bomblies & Weigel 2007), and much less is known about the evolutionary mechanisms that cause this repeatability at the phenotypic level. Moreover, in many systems it is unclear whether these common incompatibility phenotypes are developmentally similar, which can inform our understanding of the evolutionary drivers of incompatibilities. One common hybrid incompatibility in angiosperms is hybrid seed inviability (HSI).

Proper seed development is achieved through the effective coordination of three tissues with differing genetic compositions: the maternally derived seed coat, the developing embryo (1m:1p genetic composition), and a nutritive tissue that sustains the embryo until germination-the endosperm. In diploid species, seed development begins with a double fertilization event; one in which the egg cell is fertilized and the developing embryo is formed, while the other involves the fertilization of a diploid central cell that gives rise to a triploid endosperm tissue. The endosperm is thus comprised of 2m:1p genetic composition. This ratio is essential for appropriate seed development, because many genes exhibit parent-of-origin specific expression in the endosperm (i.e. imprinting; Köhler & Weinhofer-Molisch 2010). In *Arabidopsis*, this is caused primarily by DNA methylation and histone methylation (Köhler *et al*. 2012), the later of which are controlled by the Fertilization Independent Seed-Polycomb Group Complex (FIS-PcG complex; Köhler *et al*. 2003; Köhler *et al*. 2005; Gehring *et al*. 2006; Makarevich *et al*. 2006; Köhler & Makarevic 2006; Gehring *et al*. 2009; Hsieh *et al*. 2009; Wolff *et al*. 2011; Waters *et al*. 2011; Derkacheva *et al*. 2016). Indeed, imprinting in plants is thought to have evolved as a mechanism to decrease conflict over paternal and maternal interests in endosperm growth (i.e. parental conflict; Trivers 1974; Haig & Westoby 1989; Köhler *et al*. 2010).

Conflict between parents can arise in non-monogamous systems, because maternal and paternal interests in resources allocation to offspring differ (Trivers 1974; Haig & Westoby 1989). As maternity is guaranteed, selection should favor equal distribution of resources among offspring, but when paternity is variable, selection should favor the evolution of paternally derived alleles that increase resource expenditure on developing offspring (Trivers 1974; Haig & Westoby 1989). If populations have evolved differentially in this co-evolutionary arms race, then mismatches between resource-restrictive maternal alleles and resource-acquiring paternal alleles in hybrids can lead to developmental defects, and in the case of HSI, death. One prediction of parental conflict is that reciprocal F1s should show significantly different developmental trajectories, wherein one direction of the cross exhibits a paternal-excess phenotype, and the other exhibits a maternal-restrictive phenotype (Willi 2013; Raunsgard *et al*. 2018).

HSI in inter-ploidy crosses of *Arabidopsis* exhibit these two distinctive endosperm phenotypes (Scott *et al*. 1998), and endosperm defects are the cause of inviability. Both embryo and dosage rescue experiments strongly suggest that in these inter-ploidy crosses, HSI is due specifically to inappropriate development and growth of the endosperm, rather than incompatibilities arising in the embryo (Bushell *et al*. 2003; Josefsson *et al*. 2006; Dilkes *et al*. 2008; Erilova *et al*. 2009; Kradolfer *et al*. 2013). Inviability in these cases arises because of derepression of normally imprinted genes, particularly that of *ADM, SUVH7, PEG2*, and *PEG9*, which are all associated with carbohydrate sequestration and cellularization in normally developing seeds (Kradolfer *et al*. 2013; Wolff *et al*. 2015). Overexpression of these alleles causes excessive growth and improper cellularization of the endosperm associated with paternal excess, while maternal excess is often associated with precocious endosperm development (Scott *et al*. 1998; reviewed in Lafon-Placette & Köhler 2016).

While HSI is common in diploid species pairs (Tiffin *et al*. 2001), far fewer studies have determined the mechanistic underpinnings of HSI among diploids. In plants with nuclear endosperm development, such as *Arabidopsis* and *Capsella*, HSI mimic inter-ploidy crosses with a delayed or premature endosperm cellularization, depending on the paternal and maternal donor in the cross, and endosperm defects are causative of HSI (Josefsson *et al*. 2006; Burkart-Waco *et al*. 2012; Rebernig *et al*. 2015; Lafon-Placette *et al*. 2017l Lafon-Placette *et al*. 2018). In plants with cellular endosperm development (i.e. *Solanum* and *Mimulus)*, HSI is much less explored, although there is evidence to suggest that inappropriate replication of endosperm is at least partially causative of HSI in both *M. nudatus* and *M. guttatus* (Oneal *et al*. 2016), and in wild tomatoes (Lafon-Placette *et al*. 2016; Roth *et al*. 2017).

We previously reported a wide diversity of hybrid seed barriers between *M. guttatus* and its morphological variant, *M. decorus* which confer strong reproductive isolation (Coughlan *et al*. 2018). We found that *M. decorus* is multiple biological species; a diploid northern clade that exhibits strong asymmetric HSI when it is the maternal donor in crosses to *M. guttatus*; a diploid southern clade that exhibits strong asymmetric HSI when it is the paternal donor in crosses to *M. guttatus;* and at least one tetraploid taxon that exhibits variable levels of HSI (Coughlan *et al*. 2018). Because the direction of inviability differs between *M. guttatus* and each clade of diploid *M. decorus*, there must be independent evolution of maternal and paternal alleles causing inviability between northern and southern *M. decorus*, even though they are each other’s closest relative (Coughlan *et al*. 2018). HSI between *M. guttatus* and each clade of *M. decorus* thus represents two independent incidences of HSI in relatively recently diverged species. Here, we delve into the developmental basis of HSI using the *Mimulus* as a model. We test whether HSI between different diploid clades of *M. decorus* and *M. guttatus* are associated with similar developmental defects, whether these defects are endosperm related, and whether they are consistent with paternal excess/maternal restriction, as hypothesized by parental conflict. In addition, we assess the level of developmental parallelism of inviable and viable hybrid seeds between these two independent incidences of HSI.

## Methods

### Growth conditions and plant crossing

Based on previous results, *Mimulus decorus* exhibits two distinct crossing patterns: the northern clade of *M. decorus* produces inviable seeds when they are the maternal plant in crosses with *M. guttatus*, while the southern clade produces inviable seeds predominantly when they are the paternal plant in crosses with *M. guttatus*. To determine the extent of developmental parallelism, we chose a representative diploid population from each crossing type of *M. decorus* (northern clade: *IMP*; southern clade: *Odell Creek*) to cross with an annual population of *M. guttatus-* Iron Mountain (*IM*). For each population, we planted seed from 3-4 maternal families or inbred lines. For all plants, seeds were sprinkled onto moist Fafard 4-P soil, left to stratify at 4°C for 1 week, then moved to the Duke greenhouses (North Carolina, USA). Plants were transferred within the first 3 days after germination to one plant per 4” pot, then were grown in long days (18h days, 21C day/ 18C nights).

### Crossing Assays

We performed inter- and intra-specific crosses between both clades of *M. decorus* and *M. guttatus* to yield an average of six crosses for each pairwise combination of maternal families between *M. guttatus* and each clade of *M. decorus* (total of 298 crosses). Fruits were allowed to develop until ripe but before dehiscence, then seeds for each cross were counted and categorized as inviable or viable based on morphology for each fruit. Previous works suggests that morphologically assessed viability is a good proxy for germination proclivity (Garner *et al*. 2016; Coughlan *et al*. 2018).

### Microscopy and measuring growth

We sought to survey development of hybrid and parental seeds in two independent incidences of HSI. We crossed plants, as in above, but collected fruits at 4,6,8,10 and 14 Days After Pollination (DAP). Fruits were collected and immediately placed in FAA fixative (2 formaldehyde: 1 acetic acid: 10 ethanol). Plants were processed in a similar manner to White & Turner (2012). In brief, after at least 24 hours in fixative, fruits were gradually dehydrated in a Tetrabutyl-Alcohol dehydration series, then mounted in paraffin wax with ~5% gum elemi resin. Paraffin mounted specimens were then sliced to 8 micron ribbons and mounted onto slides. We performed a staining series using Safranin-O and Fast-Green, which stain for nucleic acids and carbohydrates, respectively.

We visualized and photographed at least three developing seeds from a single fruit for each cross type and time point combination (40 unique combinations, 120 seeds measured). At least 25 endosperm cell widths were measured per seed using ImageJ (Schneider *et al*. 2012).

### Statistical Analyses

Cross incompatibilities were determined with a linear mixed model using the package *lme4* and a type III ANOVA using *car* (Bates *et al*. 2015; Fox & Weisberg 2011) in the statistical interface R. All analyses were completed separately for *M. guttatus* by southern *M. decorus* and *M. guttatus* by northern *M. decorus*. For both total seed number, and proportion viable seed, we treated cross type as a fixed factor and replicate maternal and paternal lines as random effects (i.e. *Proportion viable seed ~ Cross Type + (1|maternal replicate) + (1|paternal replicate*) or *total seed count ~ Cross Type + (1|maternal replicate) + (1|paternal replicate)*). Pairwise T-tests with Holm correction for multiple testing were performed to determine which cross types varied significantly from one another. Differences in developmental trajectories were also assessed using a linear mixed model, as above. In this case, cross type, DAP, and their interaction were the fixed effects and the seed replicate was treated as a random effect.

## Results

### Post-Mating Reproductive Isolation

All lines of *M. decorus* show consistent patterns of strong, and asymmetric hybrid seed inviability when crossed with *M. guttatus*, but the two genetic clades showed oppositely asymmetric HSI (Figure 1; Odell Creek: *X^2^*= 4598, *df*=3, p<0.0001; IMP: *X^2^*= 2969, *df*=3, p<0.0001). Northern *M. decorus* produces entirely inviable seeds when the maternal donor in these crosses, while southern *M. decorus* produces entirely inviable seeds when the paternal donor (and some inviable seeds when the maternal donor; Figure 1; Coughlan *et al*. 2018). We find some evidence of pollen-pistil incompatibilities between *M. guttatus* and southern *M. decorus* (Odell Creek: X^2^= 49.33, *df*=3, p<0.0001; Figure S1), but no evidence of pollen-pistil interactions between *M. guttatus* and northern *M. decorus*, (although we note that IMP has substantially larger ovaries and produces more seeds; *X* = 23.7, *df*=3, p<0.0001; Figure S1). Inviable hybrid seeds are larger, disc-like, and shriveled, while viable hybrid seeds are smaller (as in Coughlan *et al*. 2018).

**Figure 1:**
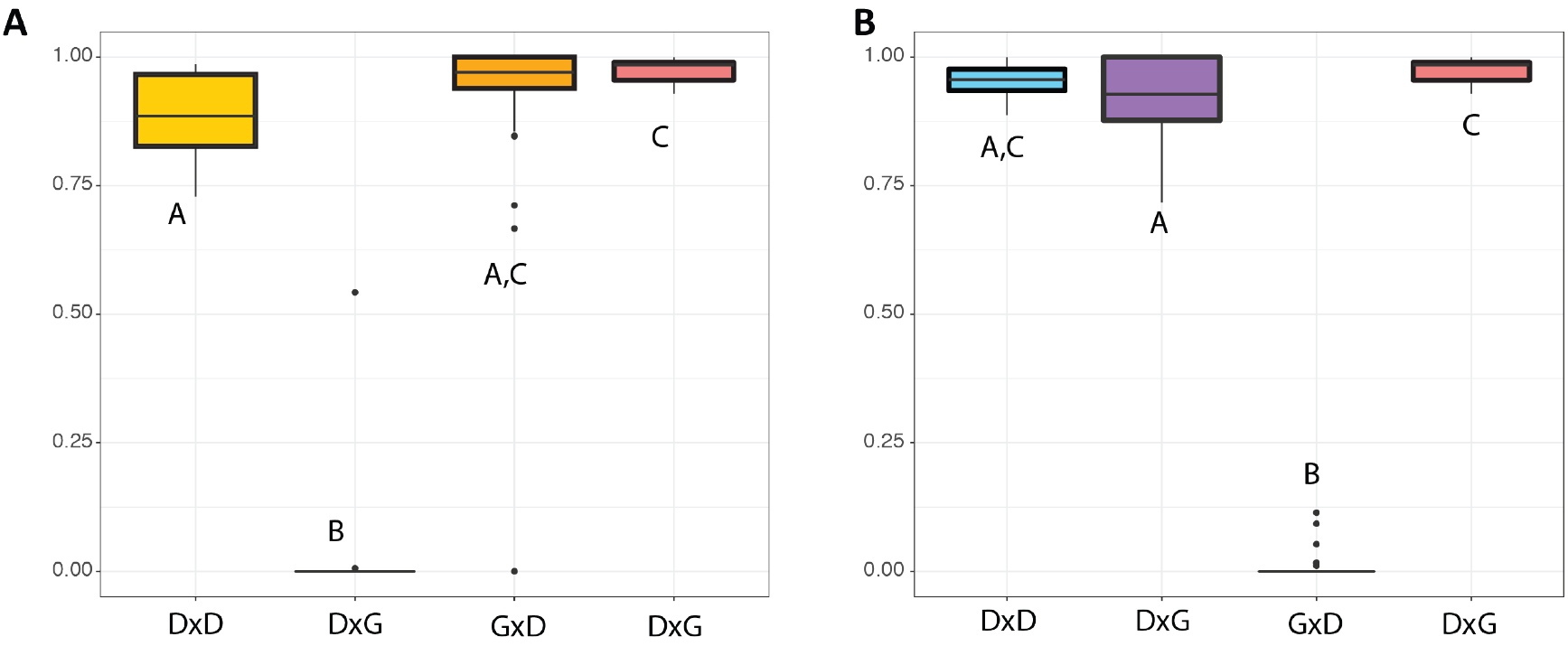
Crosses between *M. guttatus* (G) and each clade of *M. decorus* (D) shows oppositely asymmetric patterns of hybrid seed inviability. Proportion viable seed between three lines of *M. guttatus* (IM) and (A) four maternal families of northern *M. decorus* (IMP) and (B) three maternal families of southern *M. decorus* (Odell Creek). Maternal parent is listed first.

### Developmental Trajectories

The three parental species exhibit similar seed development in the context of their morphological attributes. In all three parents, endosperm is tightly packed around the embryo and continues to grow until around 8-10 DAP. At this point, the endosperm is depleted, presumably to feed the growing embryo. Embryo development among the three parents is also quite similar, with *M. guttatus* displaying a slightly accelerated growth, relative to either southern or northern *M. decorus* (i.e. *M. guttatus* reaches a heart-shaped embryo around 10 DAP, while this occurs in Odell and IMP around 14 DAP; Figure S2; Figure S3). By 14 DAP the embryo comprises most of the embryo sac, with a thin layer of endosperm remaining.

We find tremendous parent-of-origin effects on seed development in crosses between both clades *M. decorus* and *M. guttatus*. Reciprocal F1s from both sets of crosses show distinct developmental trajectories, but hybrid seeds display similar developmental trajectories, depending on seed fate (i.e. whether it will remain viable or not). In both sets of crosses, seeds that remain viable (e.g. IM×IMP and Odell×IM, with the maternal parent listed first) display a precocious development, with small endosperm cells that are quickly degraded by the embryo (Figure 2; Figure 3; Figure S2; Figure S3; Figure S4; Table S1). By 14 DAP, the only remaining endosperm is a thin layer which surrounds the embryo. Total seed size is also much smaller than either parent or the reciprocal hybrid (Coughlan *et al*. 2018). In contrast, the hybrid seeds which will eventually become inviable (i.e. IMP×IM and IM×Odell, with the maternal parent listed first) exhibit a chaotic endosperm growth, producing fewer, but larger and more diffuse endosperm cells (Figure 2; Figure 3; Figure S2; Figure S3; Table S1). Embryos in this direction of the cross remain small, relative to both parents and the reciprocal hybrid (Figure 2; Figure S2; Figure S3). By 14 DAP, most embryos were entirely degraded.

**Figure 2:**
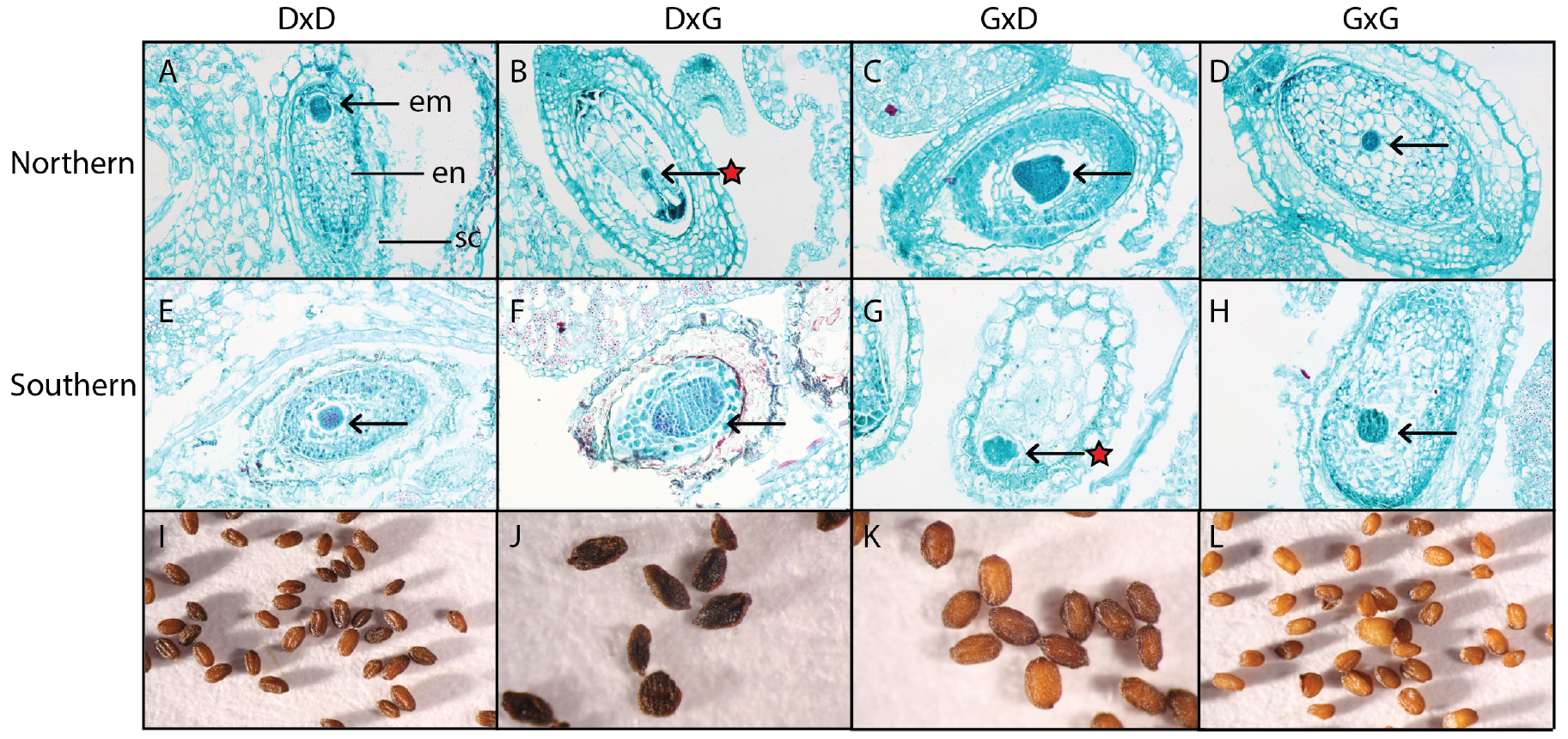
Parent of origin effects on endosperm proliferation between independent incidences of hybrid seed inviability. (A,D,E,H) Developing seeds of *M. guttatus* (‘G’) and two genetic clades of *M. decorus* (‘D’) at 8 days after pollination (DAP). Reciprocal F1s between *M. guttatus* and (B,C) the northern clade of *M. decorus* and (F,G) the southern clade of *M. decorus*. Maternal parent is listed first. Tissues are labeled in panel (A): em=embryo, en=endosperm, sc=seed coat. Arrows denote the location of the developing embryo, red stars indicate that these seeds will eventually become inviable. Fully developed seeds of (I) northern *M. decorus*, (L) *M. guttatus*, (J) a representative inviable seed, (K) a representative viable seed.

**Figure 3:**
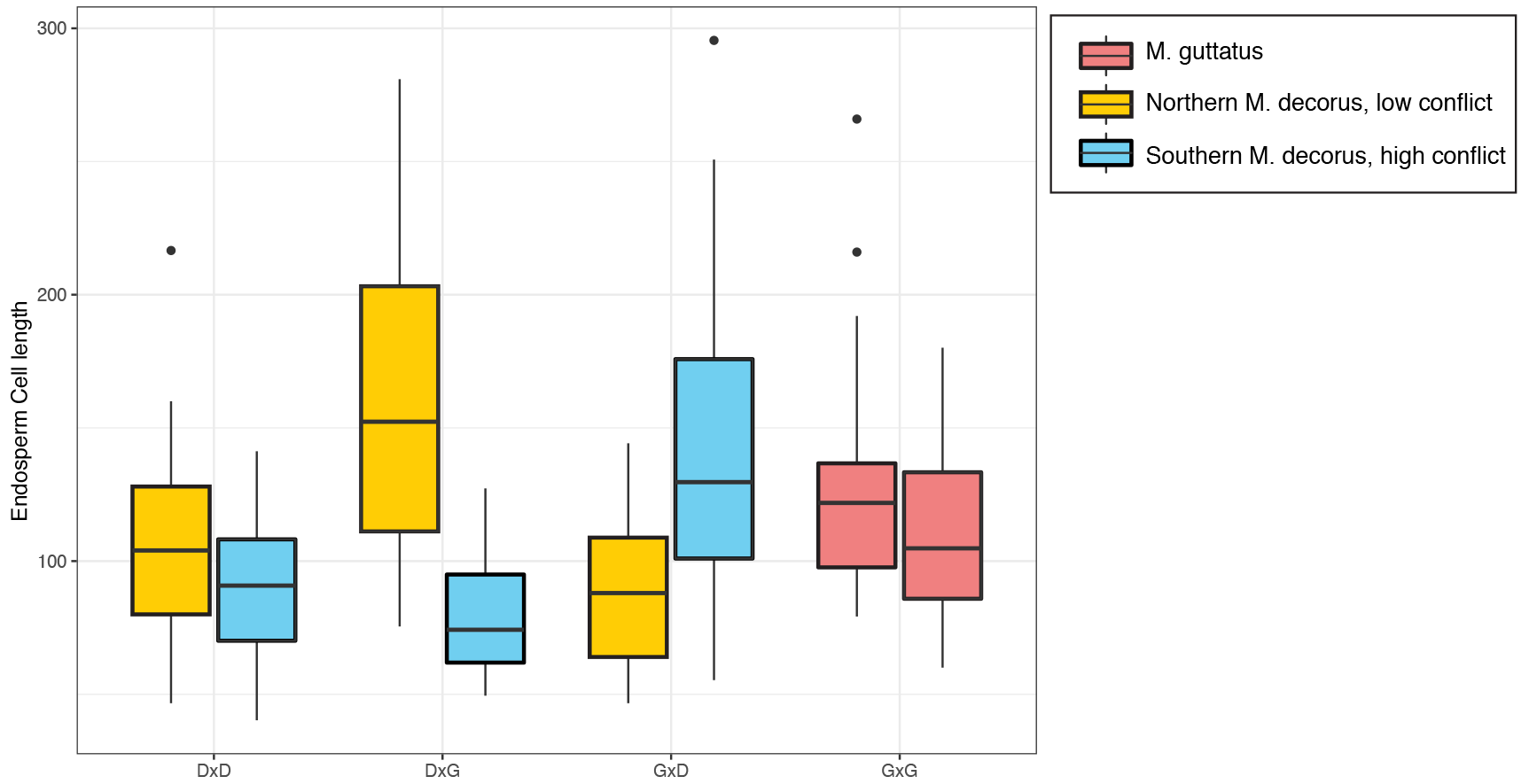
Parent-of-origin effects on endosperm growth are recapitulated at the cellular level. Cell width at 8 Days After Pollination (DAP) for each cross type (maternal parent listed first), for both *M. guttatus* crossed to IMP (yellow) and Odell Creek (blue) are shown. See Supplemental Figure 2 for all sampled time points.

While the two different sets of crosses display quite similar overall patterns of hybrid seed development, there are some notable distinctions between hybrid seeds formed between *M. guttatus* and both clades of *M. decorus*. For example, in northern *M. decorus* by *M. guttatus* crosses, where northern *M. decorus* is the maternal donor, the embryo remains comparatively small, and does not appear to grow past the 6 or 8 DAP stages. In contrast, seeds formed from between an *M. guttatus* mother and southern *M. decorus* father often exhibit a developing embryo up until 10 DAP, suggesting that the seed is aborted slightly later in this cross.

Overall, we see striking parallelism between two distinct seed incompatibilities with reciprocal F1 seeds displaying either excessive and chaotic growth patterns or precocious and stunted patterns, depending on whether the seeds are eventually aborted or not. Our findings suggest that hybrid seed inviability in this group is strongly associated with endosperm overgrowth phenotypes, though both reciprocal F1 seeds show parent-of-origin specific developmental defects.

## Discussion

The northern and southern clades of *M. decorus* exhibit strong, and asymmetric reproductive isolation with *M. guttatus*, but in opposite directions of the cross. This observation is true for all maternal family/line combinations tested here (as well as multiple populations tested in Coughlan *et al*. 2018), suggesting that HSI is not geographically restricted, as other intrinsic post-zygotic barriers have been found to be (Sweigart *et al*. 2007; Case & Willis 2008; Martin & Willis 2010; Sicard *et al*. 2015; Barnard-Kubow & Galloway 2017; Zuellig & Sweigart 2018). Total reproductive isolation in these crosses was ~52% for both species pairs, and thus may represent a significant barrier to reproduction. HSI has been shown to be a formidable barrier to reproduction in other systems (i.e. Tiffin *et al*. 2001; Lowry *et al*. 2008), but the developmental causes of this incompatibility have only been quantified for a handful of diploid species pairs. We aimed to determine the developmental trajectories of inviable and viable hybrid seeds in two independent incidences of HSI between close relatives in the *Mimulus guttatus* species complex.

We show that HSI is strongly associated with developmental defects in endosperm deposition. In inviable directions of the cross, seeds show a paternal-excess phenotype, where endosperm is chaotically distributed, and the cells are few and large. In the viable directions of the cross, seeds show a maternal-restrictive phenotype, where endosperm develops precociously, as witnessed by the general lack of endosperm remaining when seeds are fully developed, as well as the generally smaller size of both the cells and final seeds (Coughlan *et al*. 2018). Overall, these developmental patterns are consistent with a parental conflict model of hybrid inviability, wherein paternal excess is more likely to lead to seed abortion. Unfortunately, given the generally low success rate of the embryo rescues (Coughlan *unpublished*), it remains unclear whether hybrid seed inviability is due strictly to malformation of the endosperm, or whether genetic incompatibilities in the embryo also contribute.

Maternal-restriction/paternal-excess endosperm phenotypes have been shown in hybrids between wild species pairs of both *Arabidopsis* and *Capsella*, which have a nuclear-type endosperm development (Rebernig *et al*. 2015; Lafon-Placette *et al*. 2017; Lafon-Placette *et al*. 2018). In diploid *Arabidopsis* and *Capsella* crosses, as well as interploidy crosses, endosperm over-proliferation is accompanied by a failure of endosperm cells to cellularize (Scott *et al*. 1998; Bushell *et al*. 2003; Rebernig *et al*. 2015; Lafon-Placette *et al*. 2017; Lafon-Placette *et al*. 2018), and at least in *Capsella rubella* and *Capsella grandiflora*, paternal excess is associated with inviability, but maternal excess still produces viable seed (Rebernig *et al*. 2015). Findings from species with cellular-type endosperm development (e.g. wild tomatoes and other *Mimulus* species pairs) also show endosperm defects as a major cause of inviability (Roth *et al*. 2017; Oneal *et al*. 2016). However, these defects do not represent defects in rates of cellularization, but rather rates of proliferation, cell size and distribution, and optical density (Oneal *et al*. 2016; Roth *et al*. 2017). Our results are consistent with these findings, where inviable seeds display large, chaotically distributed endosperm cells, but there are fewer of them, suggesting that issues in the regular timing of cell division may be at play.

The developmental trajectories of viable versus non-viable hybrid seeds are strikingly similar between crosses of *M. guttatus* and both clades of *M. decorus*, despite the fact that inviability occurs in the opposite cross direction. This degree of developmental parallelism in independent incidences of HSI suggests that similar evolutionary processes may cause HSI within this complex. Given the maternal-restriction/paternal-excess phenotypes that we observe, one hypothesis is parental conflict. Despite considerable developmental parallelism, it remains to be seen how constrained the genetic targets of selection are between these species pairs. In other post-zygotic reproductive barriers, canonical hybrid incompatibilities often share similar genetic bases. For example, hybrid necrosis is caused by interactions involving R genes (reviewed in Bomblies & Weigel 2007), and CMS generally involves PPR genes (Polcentric Peptide Repeat genes; reviewed in Rieseberg & Blackman 2010). Whether this remains true for HSI remains to be seen. A thorough examination of the molecular and developmental genetics of HSI in this system is necessary to answer these questions, and will inform our understanding of the proximate and ultimate causes of HSI.

## Acknowledgements

We thank members of the Willis lab-particularly Elen Oneal, as well as Miguel Florez and Bob Franks for useful discussions of this project. We are grateful to Melvin Turner for teaching JMC histological techniques, and to Dick White for allowing us access to his lab space and equipment. This project was funded from NSF grants EF-0328636 and EF-0723814 to JHW, a DDIG (DEB-1501758), ASN Student research award and SSE student research award to JMC. The Duke Graduate School also provided funding to JMC with the Myra and William Waldo Boone fellowship, and financial support was provided from the Duke Biology Department.

## Data Availability

Data for the manuscript will be made publically available.

## Supplemental Figures

**Table S1:** Mixed model for type III ANOVAs for seed cell size across developmental time for IMP and Odell Creek crossed reciprocally to IM62. Cross Type= both reciprocal F1 crosses and self-fertilizations from each species treated as a separate level; treated as a fixed effect), DAP= Days After Pollination (fixed effect). Seed replicate was treated as a random effect.

|  | IMP |  |  | Odell Creek |  |  |
| --- | --- | --- | --- | --- | --- | --- |
| | $X^2$ | $DF$ | $p$ | $X^2$ | $DF$ | $p$ |
| Intercept | 295.04 | 1 | <0.0001 | 123.74 | 1 | <0.0001 |
| Cross Type | 119.848 | 3 | <0.0001 | 14.92 | 3 | 0.0018 |
| DAP | 36.033 | 1 | <0.0001 | 0.0029 | 1 | 0.96 |
| Cross Type*DAP | 42.583 | 3 | <0.0001 | 32.95 | 3 | <0.0001 |

**Figure S1:**
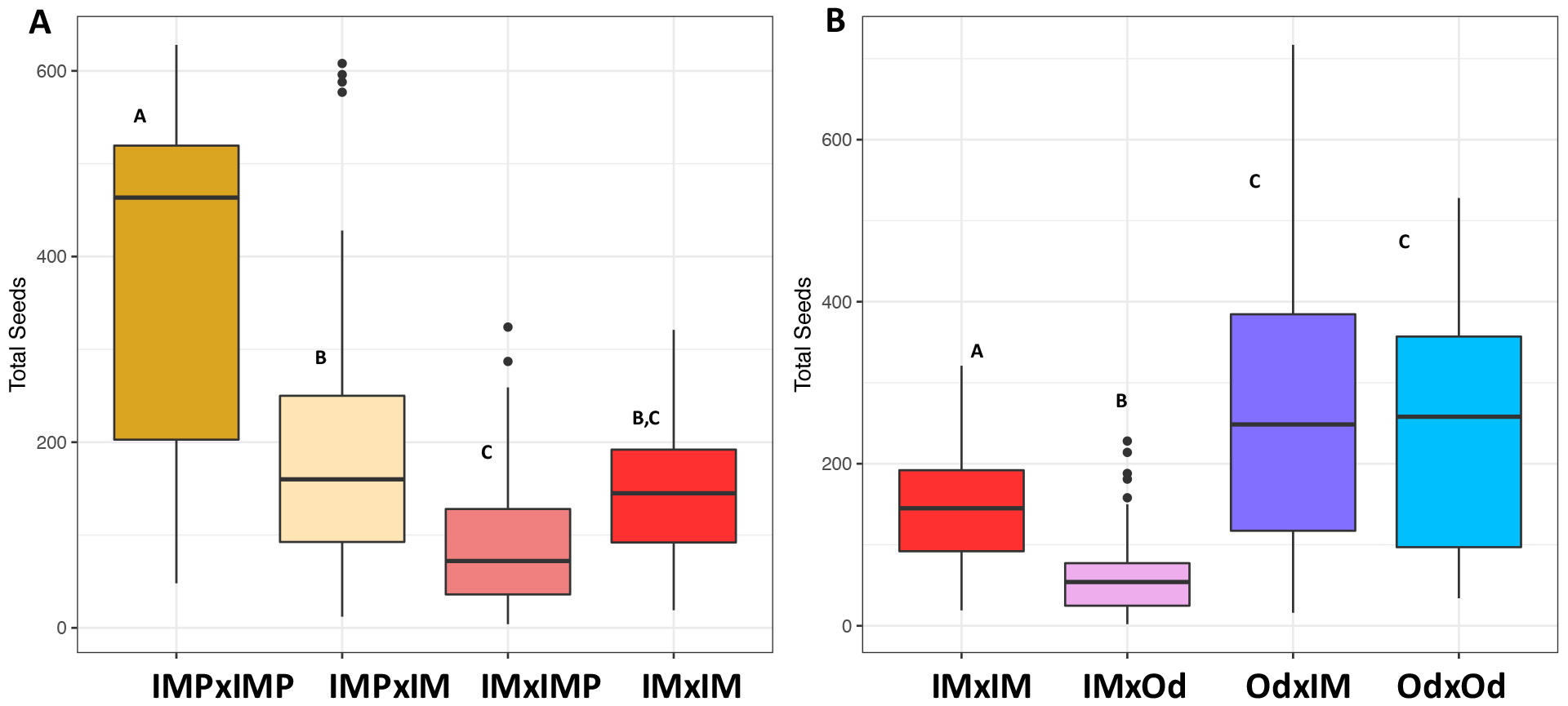
Total number of seeds in crosses for (A) three lines from IM and four maternal families of IMP selfed and crossed reciprocally and (B) three lines of IM and three maternal families of Odell Creek (Od) selfed and crossed reciprocally.

**Figure S2:**
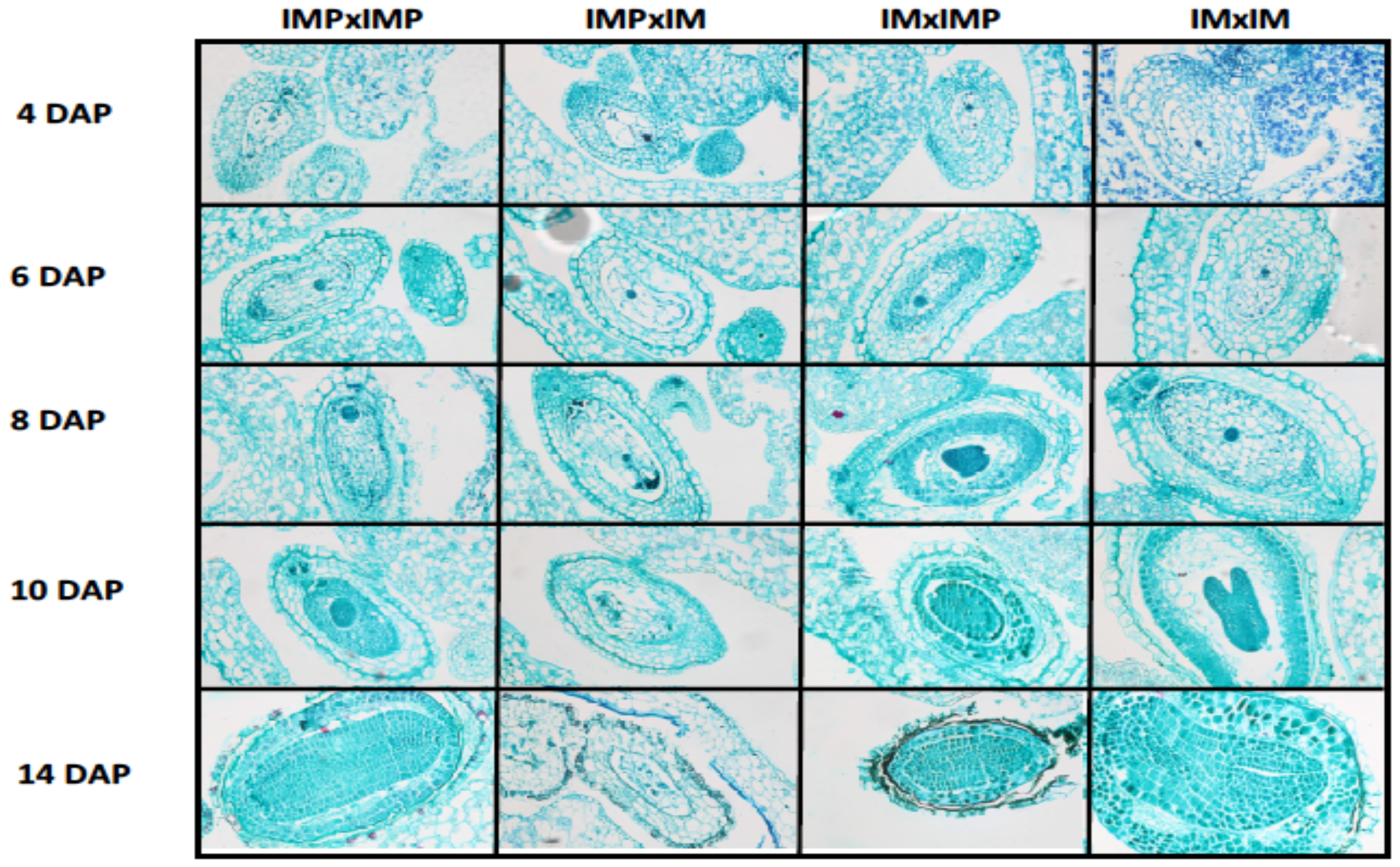
Reciprocal hybrids between *M. guttatus* and northern *M. decorus* show significant parent-of-origin effects on endosperm growth. Developing seeds between IM62 x IMP at 4, 6, 8, 10, and 14 Days After Pollination (DAP) are shown. Maternal parent is listed first.

**Figure S3:**
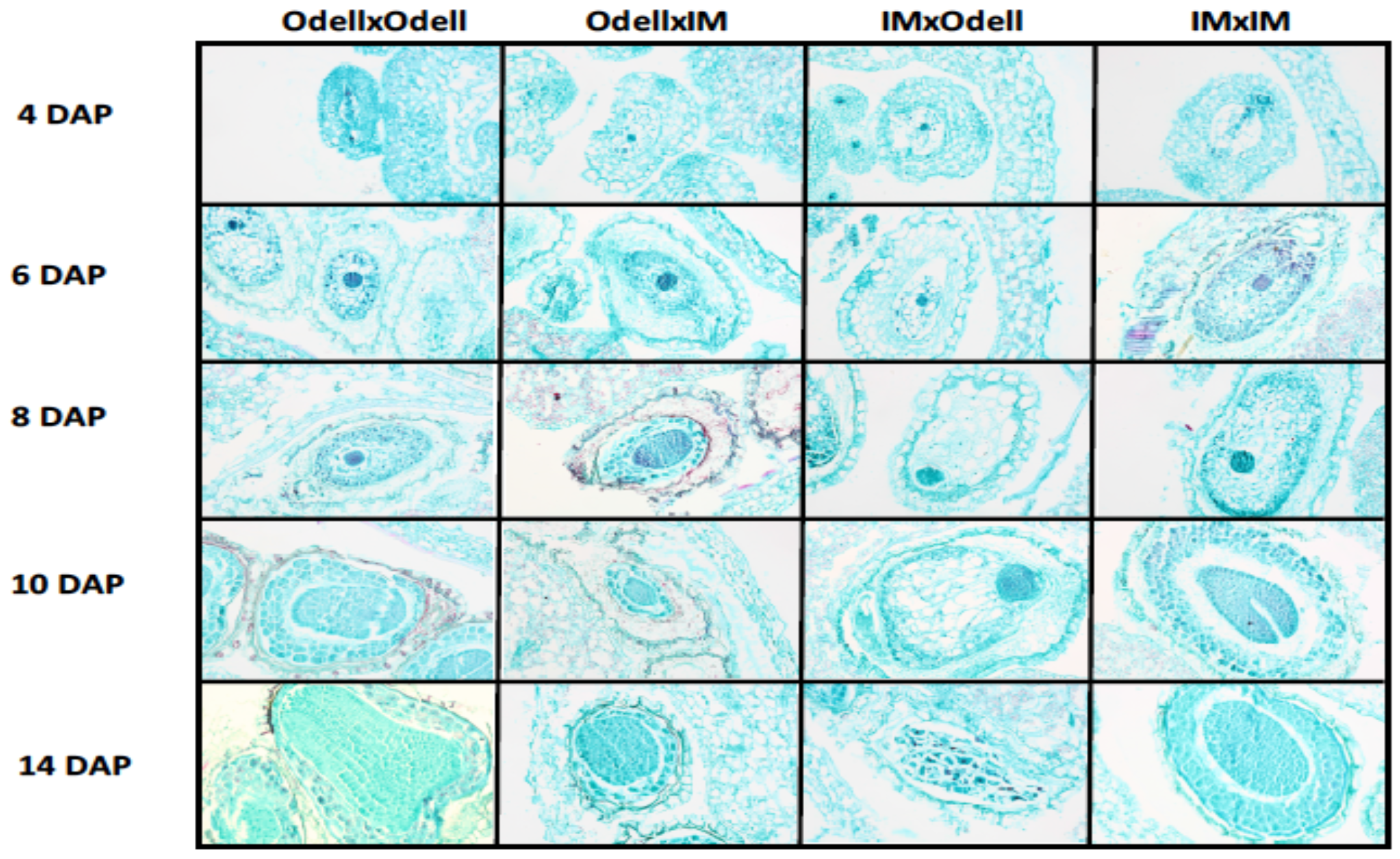
Reciprocal hybrids between *M. guttatus* and southern M. *decorus* show significant parent-of-origin effects on endosperm growth. Developing seeds between IM62 x Odell Creek at 4, 6, 8,10, and 14 Days After Pollination (DAP) are shown. Maternal parent is listed first.

**Figure S4:**
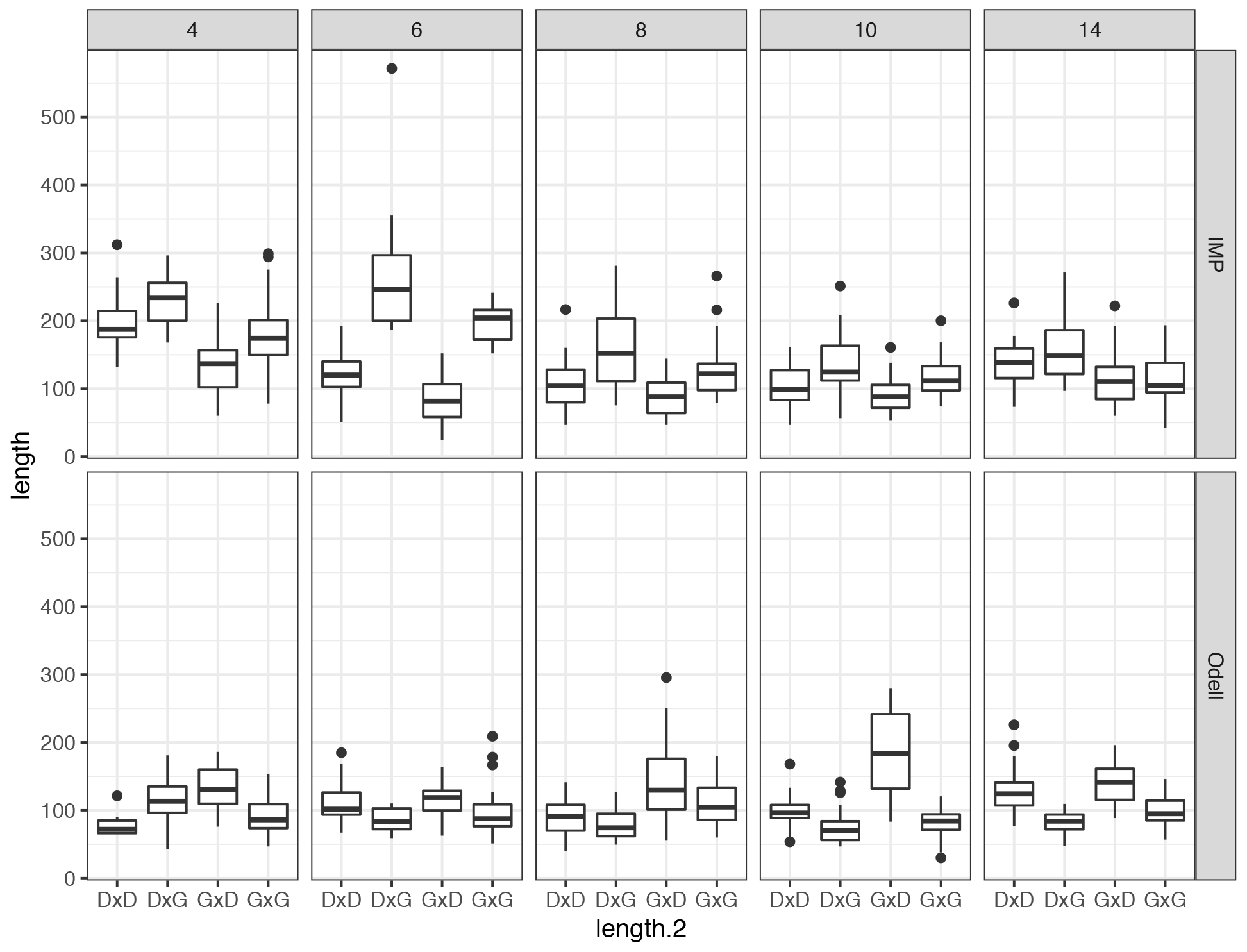
Cell lengths across all cross types for 5 developmental stages for both IMxIMP (top) and IMxOdell (bottom) crosses. Panels from left to right are each DAP.

**Author Contributions**
JMC designed this project design, performed data collection and analysis, and wrote the manuscript. JHW contributed substantially to the ideas presented here and the project design.

